# Acquisition of Host Cytosolic Protein by *Toxoplasma gondii* Bradyzoites

**DOI:** 10.1101/2020.09.11.293225

**Authors:** Geetha Kannan, Pariyamon Thaprawat, Tracey L. Schultz, Vern B. Carruthers

**Affiliations:** Department of Microbiology and Immunology, University of Michigan Medical School, Ann Arbor, MI 48109, USA

**Author notes:** Corresponding author: Vern B. Carruthers. Lycera Corp., Ann Arbor, MI 48109.

**Keywords:** chronic infection, endocytosis, parasite, protease

## Abstract

*Toxoplasma gondii* is a protozoan parasite that persists in the central nervous system as intracellular chronic stage bradyzoites that are encapsulated by a thick cyst wall. While the cyst wall separates bradyzoites from the host cytosol, it has been posited that small solutes can traverse the cyst wall to sustain bradyzoites. Recently it was found that host cytosolic macromolecules can cross the parasitophorous vacuole and are ingested and digested by actively replicating acute stage tachyzoites. However, the extent to which bradyzoites have an active ingestion pathway remained unknown. To interrogate this, we modified previously published protocols that look at tachyzoite acquisition and digestion of host proteins by measuring parasite accumulation of a host-expressed reporter protein after impairment of an endolysosomal protease (Cathepsin Protease L, CPL). Using two cystogenic parasite strains (ME49 and Pru), we demonstrate that *T. gondii* bradyzoites can ingest host-derived cytosolic mCherry. Bradyzoites acquire host mCherry within 4 hours of invasion and post-cyst wall formation. This study provides direct evidence that host macromolecules can be internalized by *T. gondii* bradyzoites across the cyst wall in infected cells.

## INTRODUCTION

*Toxoplasma gondii* is a global pathogen that impacts numerous mammalian species. Infection with this parasite can lead to blindness and encephalitis in humans as a result of reactivated infection (1, 2). The underlying source of these diseases is the chronic form of the parasite, characterized by slowly replicating bradyzoite cysts. Under specific circumstances, bradyzoites can reconvert to actively replicating tachyzoites, which induces inflammation and contributes to disease progression (3). While therapeutics exist to effectively combat tachyzoites, there is currently no fully effective treatment for bradyzoites (4). A better understanding of how bradyzoites persist is necessary to inform rationale therapeutic targeting of this elusive stage of *T. gondii*.

There is a growing cognizance of the heterogeneous and dynamic nature of bradyzoite cysts (5). During developmental switching from tachyzoite to bradyzoite, parasite proteins are secreted and localized adjacent to the parasitophorous vacuolar membrane (PVM) to form the thick glycan-rich cyst wall (6). As the cyst matures, the composition and localization of *T. gondii* proteins in the wall is altered while bradyzoites continue to replicate (7, 8). These processes require a continued source of nutrients, yet it is unclear from where bradyzoites acquire such resources and what they are.

One possibility is that host macromolecular proteins help satisfy the energy and anabolic requirements of bradyzoite cysts. The cyst wall acts as a barrier between bradyzoites and the host cytoplasm, akin to the PVM separating tachyzoites from the host cytoplasm. Yet tachyzoites can acquire across the PVM small solutes (dyes < ∼1 kDa; presumably amino acids, nucleotides, etc.) and macromolecules (GFP and mCherry) from the host cytoplasm (9-12). It has been proposed that tachyzoites ingest host cytoplasmic proteins via endocytosis at the micropore (13, 14). Micropores have also been observed in bradyzoite cysts (13). In addition, dyes (< 10 kDa) and horseradish peroxidase (HRP; 44 kDa) taken up by host cells via endocytosis have been found to enter the cyst matrix and bradyzoites within *in vitro* cysts, whereas larger proteins such as BSA (65 kDa) and transferrin (80 kDa) were observed only around the cyst wall (11, 15). Although these studies on bradyzoites suggest solutes and proteins of certain sizes that are endocytosed by host cells can be acquired by the chronic stage of *T. gondii*, they do not directly address the extent to which bradyzoites acquire proteins derived from the cytosol of infected cells.

To determine whether bradyzoites ingest host cytosolic protein, we adapted previously published protocols that demonstrated the uptake, trafficking, and accumulation of host cytosolic mCherry in tachyzoites (9, 10, 16, 17). Here we validate experimental conditions to be used to interrogate bradyzoite ingestion (e.g., parasite conversion conditions and doxycycline treatment) on tachyzoites, and provide evidence of host-derived cytosolic mCherry uptake by *T. gondii* bradyzoites.

## MATERIALS AND METHODS

### Parasite Cultures

The following strains were used in this study: ME49 wild-type, ME49 deficient CPL (MΔ*cpl*), PruΔ *ku80*SLUC (Pru), PruΔ *ku80*SLUC deficient in CPL (PΔ*cpl*). PruΔ *ku80SLUC* expresses GFP under the early bradyzoite promoter, LDH2. Details on the generation of these strains have been previously described (9, 18). *T. gondii* tachyzoites were maintained in human foreskin fibroblast (HFF) monolayers grown under standard conditions (16).

### mCherry Ingestion Assays

Parasite ingestion was determined using modifications of an already published assay (16). In brief, mCherry was expressed in inducible Chinese Hamster Ovary cells (iCHO) with the addition of 2 µg/mL of doxycycline. Parasites from infected iCHO were harvested, purified, treated with pronase and saponin, and imaged on Cell-Tak (Fisher Scientific) coated slides using a Zeiss Axiovert Observer Z1 inverted fluorescence microscope. For each biological replicate, more than 200 tachyzoites of each genotype or treatment were enumerated for host-derived mCherry accumulation within parasites. Samples were coded during the time of harvesting to blind the experimenter during imaging and quantifying.

In this study we refer to the *in vitro* conditions that promote tachyzoites conversion to bradyzoite cysts as “bradyzoite-inducing conditions”. This includes the use of alkaline (conversion) media and growth without CO_2_. For all experiments, 2 μg/mL doxycycline (DOX) was used. Detailed changes for each ingestion assay are described in the following subsections.

### Tachyzoite Ingestion under CHO vs Conversion-Inducing Conditions

mCherry was induced with DOX for 4 days in iCHO grown in standard CHO growth media (nutrient rich, 10% cosmic calf serum, Pen/Strep, pH 7.2) or conversion media (RPMI 1640 without NaHCO3, 50 mM HEPES, 5% cosmic calf serum, Pen/Strep, pH 8.3). To assess ingestion after overnight infection, iCHO were infected with tachyzoites after 4 days of induction and kept under growth and induction conditions during infection. To assess ingestion after 4 h infection, iCHO were infected after 5 days of induction and kept under growth conditions during infection. Tachyzoites were then harvested and enumerated as described above.

### Tachyzoite Ingestion with Doxycyline Treatment

The effect of doxycycline on ingestion was assessed in the following manner. Wild-type (WT, i.e., not expressing mCherry) CHO were infected with tachyzoites and grown under standard growth conditions for 5 days with DOX. In parallel, uninfected iCHO were grown under parasite conversion conditions with DOX to induce mCherry. After 5 days, tachyzoites were harvested from WT CHO and allowed to invade iCHO under conversion conditions for 4 h while kept under growth conditions. Tachyzoites were then harvested and enumerated as described above.

### Ingestion by Purified in vitro Derived Bradyzoites

Tachyzoites were converted to bradyzoite cysts in HFFs under HFF-specific conversion conditions (RPMI 1650 without NaHCO3, 50 mM HEPES, 3% FBS, Penicillin/Streptomycin, pH 8.2) for 7 days. During the last 2 days, parasites were treated with either 1 μM morpholinurea-leucine-homophenylalanine-vinyl phenyl sulfone (LHVS) or dimethyl sulfoxide (DMSO) as the solvent control. After this time, bradyzoites were harvested from *in vitro* cysts using pepsin treatment (16). Purified bradyzoites were allowed to invade iCHO under conversion conditions in the presence of DOX, LHVS (1 μM), and DMSO for either 4 h or overnight. Parasites were then harvested using saponin/pronase and the number of mCherry^+^ parasites of GFP^+^ parasites was enumerated.

### Ingestion of Purified in vivo Derived Bradyzoites

Eight-week old male CBA/J (Jackson Laboratories) mice were infected with ME49 WT tachyzoites and humanely sacrificed 5 weeks post-infection following protocols approved by the University of Michigan’s Animal Care and Use Committee. Brains from infected mice were homogenized in 1 mL of sterile Hanks Buffered Salt Solution (HBSS) and bradyzoites were subsequently harvested using pepsin treatment (19). Purified bradyzoites invaded iCHO under conversion conditions in the presence of DOX, LHVS, and DMSO for either 4 h (10 μM LHVS) or overnight (3 μM LHVS). Parasites were then harvested using saponin/pronase and the number of mCherry^+^ parasites was enumerated.

### Ingestion by in vitro Bradyzoite Cysts

Assessment of ingestion by *in vitro* bradyzoites within the cyst was done in two ways. In one set of experiments, tachyzoites were converted to bradyzoite cysts in iCHO under conversion conditions for 8-12 days. During the last 5 days of conversion, mCherry was induced with DOX. Parasites were treated with 1 μM LHVS or DMSO for the last 2 days of induction. In a second set of experiments, tachyzoites were converted to bradyzoite cysts within HFFs for 7 days. Bradyzoites were harvested via pepsin treatment and allowed to invade iCHO for 4 h before washing with media to remove extracellular bradyzoites. Bradyzoites were then kept under conversion conditions for 3 days, with the addition of DOX and 1 μM LHVS or DMSO on the last 2 days. iCHO had been treated with DOX for 2 days prior to bradyzoite infection. At the end of each experiment, bradyzoites were harvested from *in vitro* cysts with saponin/pronase. The number of mCherry^+^ parasites of GFP^+^ or BAG1^+^ parasites was enumerated.

### Immunofluorescence Staining

Purified bradyzoites were fixed in 4% paraformaldehyde, permeabilized in 0.5% Triton X-100, and stained for bradyzoite antigen 1 (BAG1) with primary rabbit anti-TgBAG1 (1:400, generated by immunization of rabbits with *E. coli*-derived recombinant BAG1) and secondary goat anti-rabbit 594 (1:1000, Invitrogen).

### Statistics

Data were analyzed using GraphPad Prism. For each data set, outliers were identified and removed using ROUT with a Q value of 0.1%. Data were then tested for normality and equal variance. If the data passed both tests, then an unpaired student’s t-test or one-way ANOVA with Dunn’s multiple comparisons was performed. If the data failed one or both tests, then a Mann-Whitney U test or Kruskal-Wallis was performed.

## RESULTS

### Parasite Conversion Conditions and Doxycycline Treatment

Tachyzoites have previously been shown to ingest host cytosolic mCherry from transiently transfected and doxycycline-induced Chinese Hamster Ovary cell lines (CHO) (16). A buildup of host-derived fluorescent protein within tachyzoites is clearly visible when parasite digestion is impaired through either genetic ablation or chemical inhibition of Cathepsin Protease L (CPL) (9, 10, 16, 17). Ingested material is observed under standard CHO culturing conditions (5% CO_2_, 10% serum, pH 7.1) and without parasite exposure to doxycycline. However, *in vitro* bradyzoites are generated under conversion-inducing conditions (ambient CO_2_, 5% serum, pH 8.3) and would be exposed to doxycycline during the induction of host mCherry. Therefore, we first sought to test whether host-cytosolic mCherry can be detected within tachyzoites that ingest under conversion-inducing conditions and after exposure to doxycycline.

To assess the extent to which parasite conversion conditions and doxycycline treatment might impair the ability of parasites to acquire host cytosolic mCherry, we performed tachyzoite ingestion assays on wild-type ME49 (WT) and genetically ablated CPL (MΔ *cpl*) tachyzoites (Fig 1). Undigested mCherry was observed within tachyzoites ingesting under conversion-inducing conditions (**Fig. 1A**). We found a significant increase in the number of mCherry positive MΔ *cpl* tachyzoites compared to WT under conversion-inducing conditions (**Fig. 1B**). This increase was observed in MΔ *cpl* at 4 h post-infection and after overnight replication of parasites in mCherry expressing iCHO. The same results were observed under standard CHO culturing conditions, which served as a positive control for the assay (**Fig. 1C**). In addition, there was an increase in the number of mCherry positive MΔ *cpl* compared to WT tachyzoites that were pre-treated with doxycycline (+ Dox) for 5 days, which was also observed in untreated (- DOX) controls (Fig. 1D). Taken together, these findings indicate that host cytosolic mCherry can be ingested by tachyzoites under conversion-inducing culture conditions and with exposure to doxycycline.

**Figure 1:**
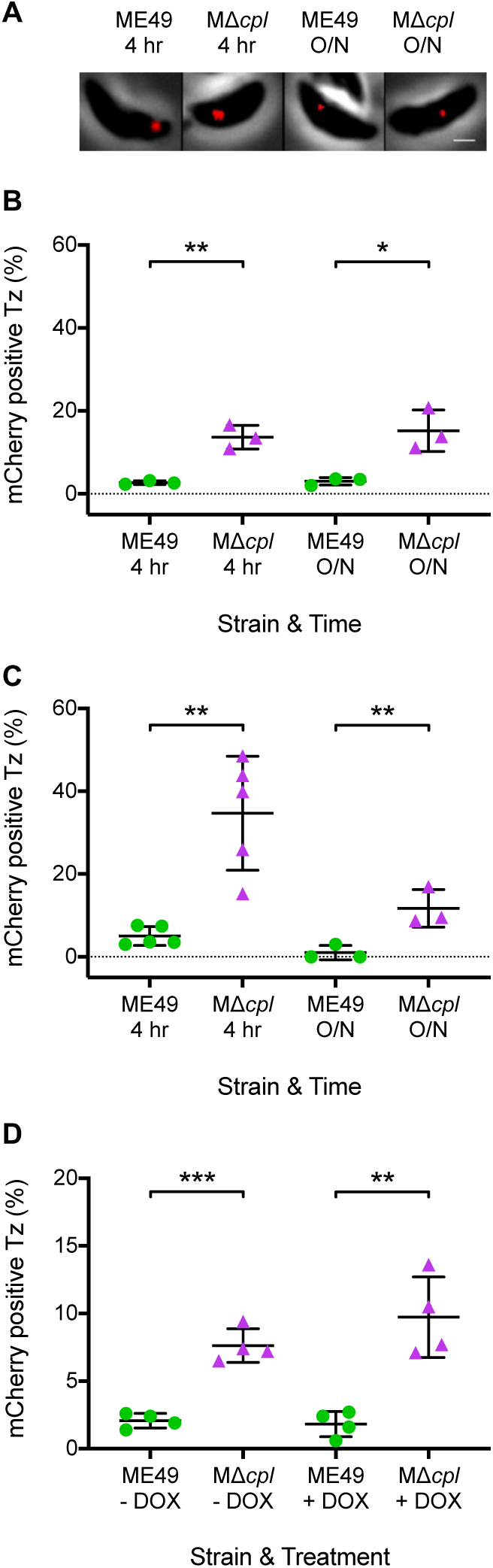
Ingestion of host cytosolic material by tachyzoites under bradyzoite-inducing conditions and doxycycline treatment. A. Representative images of tachyzoites with ingested host-cytosolic mCherry under bradyzoite-inducing conditions. Scale bar denotes 1 µM. B. Ingestion of host cytosolic mCherry by *T. gondii* tachyzoites under bradyzoite-inducing conditions. Three independent experiments were performed for each strain. The following number of parasites were enumerated for each experiment. ME49 4 h (280, 265, 256), MΔ *cpl* (304, 271, 303), ME49 O/N (199, 228, 220), MΔ *cpl* O/N (236, 195, 287). Unpaired t-test was performed to compare genotypes within each time point. * denotes p < 0.05 and ** denotes p < 0.005. C. Ingestion of host cytosolic mCherry by *T. gondii* tachyzoites under CHO culturing conditions. Three or five independent experiments were performed. The following number of parasites were enumerated for each experiment. ME49 4 h (281, 227, 259, 244, 262), MΔ *cpl* (163, 228, 263, 283, 272), ME49 overnight (O/N) (194, 180, 220, 340, 293, 529), MΔ *cpl* O/N (218, 295, 231, 328, 238, 322). Mann-Whitney U-test was performed to compare genotypes within each time point. * denotes p < 0.05 and ** denotes p < 0.005. D. Doxycycline does not affect ingestion of host cytosolic mCherry by *T. gondii* tachyzoites. Four independent experiments were performed. The following number of parasites were enumerated for each experiment. ME49 – DOX (318, 271, 258, 210), MΔ *cpl* – DOX (214, 277, 230, 292), ME49 + DOX (226, 338, 248, 206), MΔ *cpl* + DOX (324, 235, 222, 266). Unpaired t-test was performed to compare strains within each treatment. ** denotes p<0.005 and *** denotes p<0.0005.

### Bradyzoite Ingestion of Host-Derived mCherry

We next wanted to determine whether bradyzoites can ingest host cytosolic mCherry. To do this we performed the ingestion assay with either *in vivo-* or *in vitro*-derived bradyzoites that were liberated from cysts (**Fig. 2**). Because an insufficient number of MΔ *cpl* cysts can be recovered from the rodent brain (18), we used the irreversible CPL inhibitor, morpholinurea-leucine-homophenylalanine-vinyl phenyl sulfone (LHVS), to limit parasite digestion during and post-invasion. As with MΔ *cpl* tachyzoites (**Fig 1A**), a significantly greater percentage of LHVS-treated *in vivo-*derived ME49 bradyzoites contained host mCherry when compared to the DMSO control (**Fig. 2A**). This was seen both at 4 h post-infection and after overnight replication. We found the same results with *in vitro-*derived bradyzoites of a separate cystogenic strain, Pru (**Fig. 2B**). Taken together, these data suggest that *T. gondii* bradyzoites can acquire host cytosolic mCherry within 24 h of invasion.

**Figure 2:**
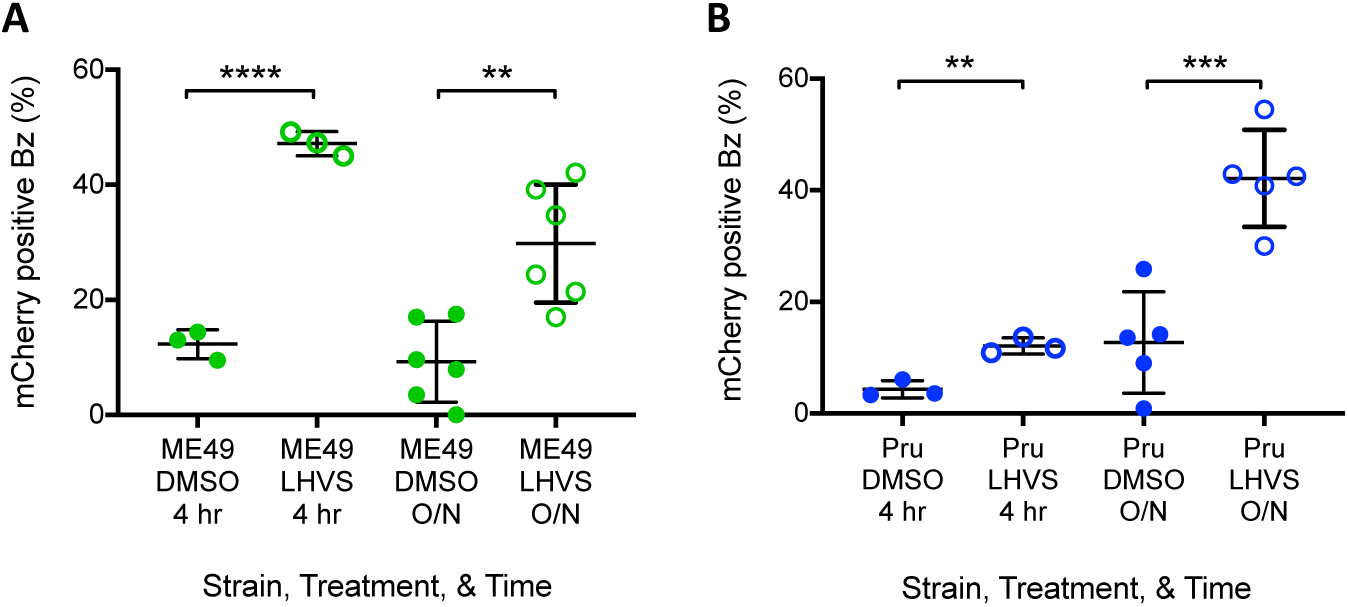
Bradyzoites can ingest host-derived mCherry. A. Bradyzoites from mouse brain cysts can ingest host cytosolic mCherry. Three to six independent experiments were performed. The following number of parasites were enumerated for each experiment. 4 h: ME49 DMSO (236, 210, 277), ME49 LHVS (10 µM) (255, 299, 245); overnight (O/N): ME49 DMSO (282, 229, 217, 245, 228, 245), ME49 LHVS (3 µM) (266, 394, 234, 245, 202, 217). Unpaired t-test was performed to compare treatments within each time point. ** denotes p< 0.005 and **** denotes p< 0.0001. B. Ingestion of host cytosolic mCherry by *in vitro* bradyzoites. Three to five independent experiments were performed. The following number of GFP+ parasites were enumerated for each experiment. 4 hr: Pru DMSO (365, 231, 214), Pru LHVS (1 µM) (213, 238, 175); O/N: Pru DMSO (231, 189, 154, 212, 231), Pru LHVS (1 µM) (318, 245, 213, 200, 242). Unpaired t-test was performed to compare treatments within each time point. ** denotes p < 0.005 and *** denotes p < 0.0005.

### Ingestion of mCherry After Cyst Wall Formation

We next wanted to determine whether bradyzoites can acquire host-derived mCherry through the cyst wall (**Fig. 3**). To elucidate this, we infected iCHO with ME49 or Pru tachyzoites and converted them to bradyzoites for 7 days prior to doxycycline induction for 5 days. There was a significant increase in host-derived mCherry upon chemical and genetic ablation of CPL in ME49 (**Fig. 3A**) and Pru (**Fig. 3B**) bradyzoites as compared with WT DMSO controls. The same results were obtained when Pru parasites were converted for a shorter time (**Fig. 3C**). However, when in culture for nearly a week, *in vitro* bradyzoites can egress out of a cyst to invade adjacent host cells. Because the bradyzoites here were in culture for 8-12 days, it is possible the ingested mCherry we observe is from newly invaded parasites rather than by passing through the cyst wall. To address this, we shortened the time of conversion within iCHO by starting with purified *in vitro* derived bradyzoites and washed off extracellular bradyzoites after a 4 h invasion. At 24 h post-infection, when the cyst wall has already started forming around replicated bradyzoites (7), LHVS treatment began for 2 days. Because CPL is not inhibited until after the wall begins to form, it is more likely that any mCherry that accumulates within the parasite must have done so through the cyst wall. Indeed, we found a greater percentage of LHVS-treated ME49 and Pru parasites with mCherry than DMSO controls (**Fig. 3D**). A schematic for this experiment is provided in **Fig. 3E**. Taken together, this data suggests that *T. gondii* bradyzoites can acquire host cytosolic protein through the cyst wall.

**Figure 3:**
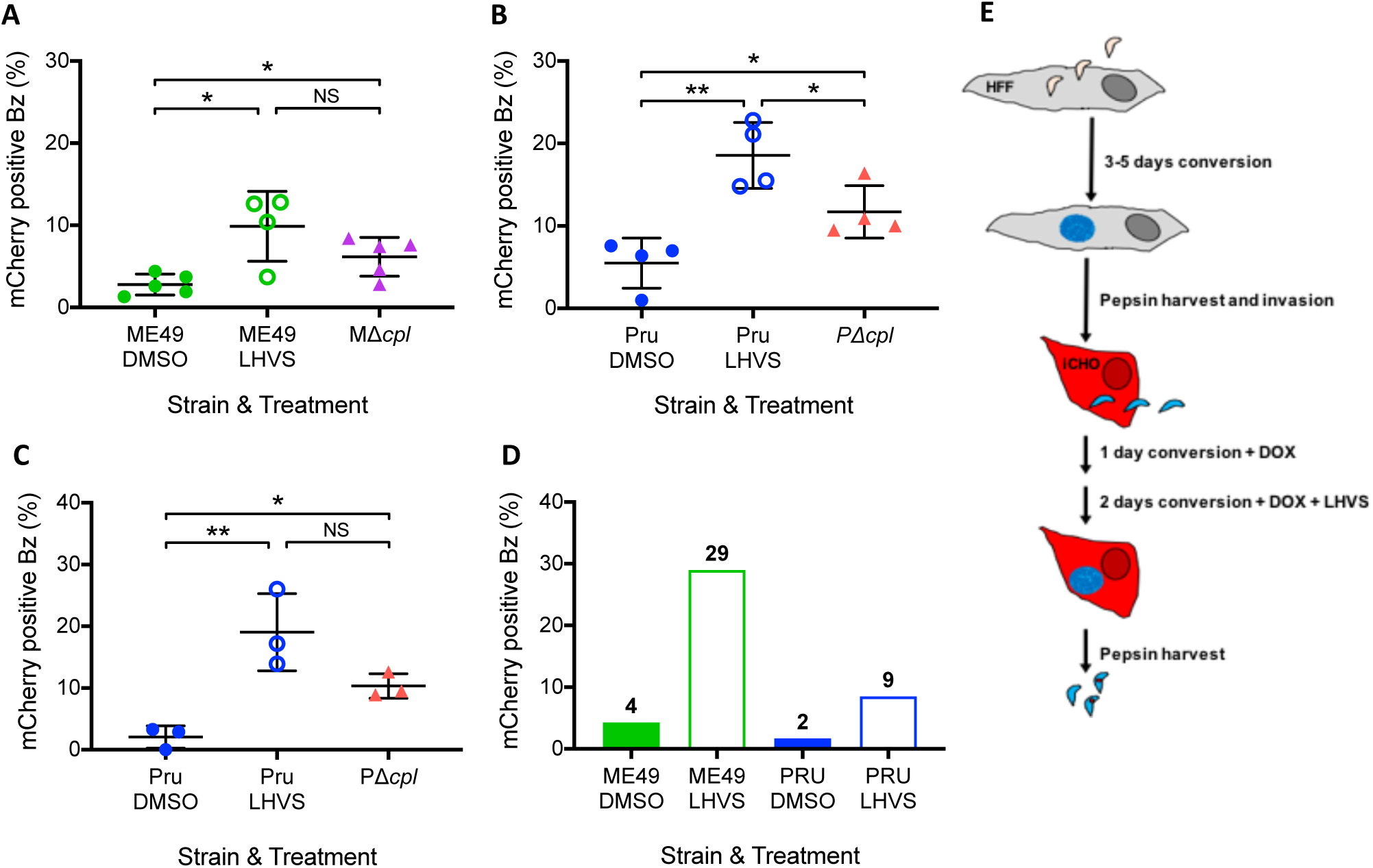
*in vitro* bradyzoite cysts contain host-derived mCherry. A. Ingestion of host cytosolic mCherry by *T. gondii in vitro* bradyzoites converted in iCHO for 12 days and treated with LHVS for 2 days, BAG1 + stained parasites were enumerated. 4-5 independent experiments were performed for each strain and treatment. The following number of parasites were enumerated for each experiment. ME49 DMSO (262, 342, 272, 237, 272), ME49 LHVS (231, 195, 328, 231), MΔ *cpl* (95, 254, 282, 135, 251). Mann-Whitney U test was used to compare groups. * denotes p< 0.05 and ** denotes p < 0.005. B. Ingestion of host cytosolic mCherry by *T. gondii in vitro* bradyzoites converted in iCHO for 12 days and treated with LHVS for 2 days; GFP+ parasites were enumerated. Four independent experiments were performed for each strain and treatment. The following number of parasites were enumerated for each experiment. Pru DMSO (85, 277, 264, 298), Pru LHVS (209, 215, 223, 317), PΔ *cpl* (270, 432, 232, 276). Unpaired t-test was used to compare all groups. * denotes p< 0.05 and ** denotes p < 0.005. C. Ingestion of host cytosolic mCherry by *T. gondii in vitro* bradyzoites converted in iCHO for 8 days and treated with LHVS for 2 days; GFP+ parasites were enumerated. 3 independent experiments were performed for each strain and treatment. The following number of parasites were enumerated for each experiment. Pru DMSO (270, 210, 209), Pru LHVS (357, 227, 209), PΔ *cpl* (464, 198, 224). Unpaired t-test was used to compare all groups. * denotes p< 0.05 and ** denotes p < 0.005. D. Ingestion of host cytosolic mCherry by *T. gondii in vitro* bradyzoites converted in HFFs for 7 days, harvested, and then converted in iCHO for 3 days with 1 µM LHVS for 2 days; BAG1+ ME49 and GFP+ Pru bradyzoites were enumerated. The numbers above each bar represent the % mCherry positive bradyzoites from 1 experiment. The following number of parasites were analyzed in each group: ME49 DMSO (234), ME49 LHVS (281), Pru DMSO (230), Pru LHVS (212). E. Schematic of experimental design for D.

## DISCUSSION

Here we provide evidence that *Toxoplasma* bradyzoites can ingest host-cytosolic mCherry. This acquisition of host protein by bradyzoites occurs within 4 h of invasion as well as after 24 h of bradyzoite infection, when the cyst wall would have started forming (7). Our findings are therefore consistent with host proteins potentially serving as a nutrition source for bradyzoite cysts.

Our study builds upon prior discoveries that the cyst wall allows the passing of large dyes (<10 kDa) and HRP (44 kDa) that can get incorporated into bradyzoites (11, 15). Utilizing tracers of different sizes for bulk and receptor-mediated endocytosis revealed that bradyzoites may only incorporate proteins up to 44 kDa via bulk endocytosis (15). However, these tracers are taken up by the host cell from the extracellular environment through endocytosis, trafficked to the host lysosomes that decorate the outside of the cyst wall, and then material up to a certain size is taken into the cyst. It is uncertain whether the host-endocytosed material enters bradyzoites via bulk or receptor-mediated endocytosis. Nevertheless, in contrast to these studies, ours shows another source of resources for bradyzoites: host-derived cytosolic proteins.

There are two limitations to our study. The first consideration is that host-cytosolic mCherry within bradyzoites may have been acquired in tachyzoite-like parasites (e.g., not fully mature bradyzoites). Indeed, it has been shown in studies that tachyzoites are capable of ingesting host-cytosolic mCherry (10, 16, 17). Although tachyzoites are killed during the pepsin harvesting of bradyzoites from cysts, we cannot rule out the possibility that partially-converted tachyzoites within *in vitro* cysts survived harvesting and were the ones to ingest host material, or that once added to culture, fully mature bradyzoites from the mouse brain did not begin expressing tachyzoite proteins that would aid in ingestion. Live cell imaging with parasites expressing stage specific fluorescent proteins and host cells that express cytoplasmic proteins that only fluoresce once inside the parasite lysosome would aid in clearly demonstrating that bradyzoites are capable of ingesting host cytosolic protein.

The second limitation is that we do not know the maturity of *in vitro* cysts within iCHO. It has recently been shown that the architecture and composition of the cyst wall changes during maturation (7, 8). While we found host-derived mCherry within bradyzoites that had CPL inhibited after the cyst wall would have started forming, it is conceivable that the wall mirrored that of an immature cyst. It is therefore unclear whether host-derived proteins would be capable of passing through a fully mature cyst wall, as would be found *in vivo*. Staining and analysis of the localization of different proteins within the cyst wall could discern the maturity of *in vitro* cysts and at what level of development the parasites may stop utilizing host-cytosolic proteins for nutrients and instead move to an alternative energy source.

In order for *T. gondii* to survive and thrive, the parasite acquires a multitude of nutrients from the host cell via various mechanisms. For instance, tachyzoites obtain host-derived fatty acids, proteins, and small molecules/amino acids to support replication. While host-derived protein can be acquired within minutes of parasite invasion and after the PVM has formed, small molecules and amino acids are obtained after PVM formation through the GRA17/23 putative nutrient pore, and fatty acids traverse the PVM during tachyzoite replication (9, 12, 20). It would be unsurprising that with their newly appreciated dynamic qualities, bradyzoite cysts also persist by obtaining nutrients from the host cell through different sources. For example, *in vitro T. gondii* bradyzoites form more lipid droplets when oleic acid (OA) is added to the culture media, suggesting parasite acquisition of host-derived fatty acids (21). More recently, GRA17 has also been posited to provide nutrients to bradyzoite cysts during differentiation, suggested by the decreased viability in ME49Δ*gra17* parasites (22). Our current study adds host-cytoplasmic proteins to the list of host-derived components that bradyzoites can acquire.

To better understand the importance of host-cytosolic protein acquisition by *T. gondii* bradyzoites for parasite persistence, more experiments are needed to gain clarity and mechanistic insight. As mentioned in the second caveat to this study, it would be important to have an understanding as to the point in cyst maturation that host-derived proteins are a key source of nutrients as compared to the other sources. In addition, it is necessary to elucidate a mechanism by which *T. gondii* bradyzoites uptake host cytoplasmic protein. Inhibiting acquisition of host-derived protein uptake would enable studies to determine the importance of this nutrient pathway in the viability and persistence of *T. gondii* bradyzoites.

## AUTHOR CONTRIBUTIONS

GK designed, performed, and analyzed experiments and wrote the first manuscript draft. PT performed and analyzed experiments. TS supplied *in vivo* cysts and provided technical assistance. VBC conceived the study and revised the manuscript.

## ACKNOWLEDGEMENTS

This work was supported by National Institutes of Health grant R01AI120627 (VBC). The authors appreciate feedback on manuscript drafts from Drs. My-Hang Huynh and Zhicheng Dou. We thank Dr. Louis Weiss for providing the plasmid for expression of BAG1 in *E. coli*.

## REFERENCES

1. Winstanley P. 1995. Drug treatment of toxoplasmic encephalitis in acquired immunodeficiency syndrome. Postgrad Med J 71:404–8.

2. Commodaro AG, Belfort RN, Rizzo LV, Muccioli C, Silveira C, Burnier MN, Jr., Belfort R, Jr. 2009. Ocular toxoplasmosis: an update and review of the literature. Mem Inst Oswaldo Cruz 104:345–50.

3. Lyons RE, McLeod R, Roberts CW. 2002. Toxoplasma gondii tachyzoite-bradyzoite interconversion. Trends Parasitol 18:198–201.

4. Konstantinovic N, Guegan H, Stajner T, Belaz S, Robert-Gangneux F. 2019. Treatment of toxoplasmosis: Current options and future perspectives. Food Waterborne Parasitol 15:e00036.

5. Watts E, Zhao Y, Dhara A, Eller B, Patwardhan A, Sinai AP. 2015. Novel Approaches Reveal that Toxoplasma gondii Bradyzoites within Tissue Cysts Are Dynamic and Replicating Entities In Vivo. mBio 6:e01155–15.

6. Dubey JP, Lindsay DS, Speer CA. 1998. Structures of Toxoplasma gondii tachyzoites, bradyzoites, and sporozoites and biology and development of tissue cysts. Clin Microbiol Rev 11:267–99.

7. Guevara RB, Fox BA, Bzik DJ. 2020. Toxoplasma gondii Parasitophorous Vacuole Membrane-Associated Dense Granule Proteins Regulate Maturation of the Cyst Wall. mSphere 5.

8. Guevara RB, Fox BA, Falla A, Bzik DJ. 2019. Toxoplasma gondii Intravacuolar-Network-Associated Dense Granule Proteins Regulate Maturation of the Cyst Matrix and Cyst Wall. mSphere 4.

9. Dou Z, McGovern OL, Di Cristina M, Carruthers VB. 2014. Toxoplasma gondii ingests and digests host cytosolic proteins. MBio 5:e01188–14.

10. McGovern OL, Rivera-Cuevas Y, Kannan G, Narwold AJ, Jr., Carruthers VB. 2018. Intersection of endocytic and exocytic systems in Toxoplasma gondii. Traffic 19:336–353.

11. Lemgruber L, Lupetti P, Martins-Duarte ES, De Souza W, Vommaro RC. 2011. The organization of the wall filaments and characterization of the matrix structures of Toxoplasma gondii cyst form. Cell Microbiol 13:1920–32.

12. Gold DA, Kaplan AD, Lis A, Bett GC, Rosowski EE, Cirelli KM, Bougdour A, Sidik SM, Beck JR, Lourido S, Egea PF, Bradley PJ, Hakimi MA, Rasmusson RL, Saeij JP. 2015. The Toxoplasma Dense Granule Proteins GRA17 and GRA23 Mediate the Movement of Small Molecules between the Host and the Parasitophorous Vacuole. Cell Host Microbe 17:642–52.

13. Nichols BA, Chiappino ML, Pavesio CE. 1994. Endocytosis at the micropore of Toxoplasma gondii. Parasitol Res 80:91–8.

14. Spielmann T, Gras S, Sabitzki R, Meissner M. 2020. Endocytosis in Plasmodium and Toxoplasma Parasites. Trends Parasitol 36:520–532.

15. Paredes-Santos TC, Martins-Duarte ES, de Souza W, Attias M, Vommaro RC. 2018. Toxoplasma gondii reorganizes the host cell architecture during spontaneous cyst formation in vitro. Parasitology 145:1027–1038.

16. Kannan G, Di Cristina M, Schultz AJ, Huynh MH, Wang F, Schultz TL, Lunghi M, Coppens I, Carruthers VB. 2019. Role of Toxoplasma gondii Chloroquine Resistance Transporter in Bradyzoite Viability and Digestive Vacuole Maintenance. mBio 10.

17. McDonald C, Smith D, Di Cristina M, Kannan G, Dou Z, Carruthers VB. 2020. Toxoplasma Cathepsin Protease B and Aspartyl Protease 1 Are Dispensable for Endolysosomal Protein Digestion. mSphere 5.

18. Di Cristina M, Dou Z, Lunghi M, Kannan G, Huynh MH, McGovern OL, Schultz TL, Schultz AJ, Miller AJ, Hayes BM, van der Linden W, Emiliani C, Bogyo M, Besteiro S, Coppens I, Carruthers VB. 2017. Toxoplasma depends on lysosomal consumption of autophagosomes for persistent infection. Nat Microbiol 2:17096.

19. Mayoral J, Di Cristina M, Carruthers VB, Weiss LM. 2020. Toxoplasma gondii: Bradyzoite Differentiation In Vitro and In Vivo. Methods Mol Biol 2071:269–282.

20. Nolan SJ, Romano JD, Coppens I. 2017. Host lipid droplets: An important source of lipids salvaged by the intracellular parasite Toxoplasma gondii. PLoS Pathog 13:e1006362.

21. Nolan SJ, Romano JD, Kline JT, Coppens I. 2018. Novel Approaches To Kill Toxoplasma gondii by Exploiting the Uncontrolled Uptake of Unsaturated Fatty Acids and Vulnerability to Lipid Storage Inhibition of the Parasite. Antimicrob Agents Chemother 62.

22. Paredes-Santos T, Wang Y, Waldman B, Lourido S, Saeij JP. 2019. The GRA17 Parasitophorous Vacuole Membrane Permeability Pore Contributes to Bradyzoite Viability. Front Cell Infect Microbiol 9:321.

